# The Relative Roles of Voice and Gesture in Early Communication Development

**DOI:** 10.1101/2020.12.07.415232

**Authors:** Megan M. Burkhardt-Reed, Helen L. Long, Dale D. Bowman, Edina R. Bene, D. Kimbrough Oller

**Affiliations:** School of Communication Sciences and Disorders, University of Memphis, Memphis, TN, USA; Department of Mathematics, University of Memphis, Memphis, TN, USA; Institute for Intelligent Systems, University of Memphis, Memphis, TN, USA; Konrad Lorenz Institute for Evolution and Cognition Research, Klosterneuburg, Austria

**Keywords:** Infant gesture, infant vocalization, prelinguistic communication, language development, origins of language

## Abstract

Both vocalization and gesture are universal modes of communication and fundamental features of language development. Many believe that language evolved out of early gestural use; however, evidence reported here suggests vocalization precedes gesture in human communication and forms the predominant foundation for language. To our knowledge no prior research has investigated the rates of emergence of both gesture and vocalization in human infants to evaluate this question. We evaluated the rates of gesture and speech-like vocalizations (protophones) of 10 infants at 4, 7, and 11 months of age using parent-infant laboratory recordings. We found that infant protophones outnumbered gestures substantially at all three ages, ranging from >30 times more protophones than gestures at 3 months, to more than twice as many protophones as gestures at 11 months. The results suggest that vocalization is the predominant mode of communication in human infants from the beginning of life.

## 1. Introduction

The origin of language has been a longstanding focus of speculation and investigation (De Condillac, 1756; Hewes, 1973; Corballis, 2010). Theoretical debates about the origin of language center on whether vocalization or gesture played a more fundamental role, with prominent thinkers supporting both possibilities (Call & Tomasello, 2007; Armstrong & Wilcox, 2007; Liszkowski et al., 2011; Sterelny, 2012; Gillespie-Lynch et al., 2014; Oller et al., 2019). Both vocal and gestural theories commonly reference the communicative behaviors of nonhuman primates, primarily the great apes, as evidence regarding the origin of language. Since great apes are claimed to communicate more flexibly in a visual-gestural modality than a vocal one, it has often been argued that our early hominin ancestors must have been gestural communicators.

In support of a vocal source, some suggest that the vocal capabilities of the great apes have been largely underestimated (Cheney & Seyfarth, 2005; Lameira, 2017). Also, it has been noted that vocalization has the advantage of being able to function when the communicators are in darkness or are not looking at each other (Kendon, 2017). The massive amount of early infant vocal behavior and communication has also been emphasized by those who advocate a primarily vocal foundation for language. It has been claimed that the endogenous nature of early human infant vocal behaviors provides evidence for a foundational role of vocalization in language acquisition (Iyer et al., 2016; Oller et al., 2019, Long et al., 2020). However, the widespread claim that gesture paves the way for language development in modern human infants has furthered speculation about gestural origins (Iverson & Goldin-Meadow, 2005; Gillespie-Lynch et al., 2013). Research on prelinguistic communicative behaviors suggests that both vocalization and gesture are fundamental features in language development. The present papers aims to quantify the rates of speech-like vocalization and gesture across the first year of life to gain insight on which modality may play a more foundational role. We also address the extent to which gestures and vocalizations tend to be directed to potential receivers. We begin by evaluating empirical reports on both gesture and vocalization in human development.

### 1.1. Gesture as a Foundational Activity in Language

Gesture is widely viewed as a key foundational activity in the emergence of spoken language. Although infants vocalize from birth, many believe gesture is the first opportunity for prelinguistic infants to convey messages that they have yet to gain the words to express (Bates, 1975; Iverson & Goldin-Meadow, 2005; Silva Lima & Cruz-Santos, 2012). Several research studies have presented evidence that suggests gestures are the primary driving force for the development of symbols (Bates et al., 1979, Goodwyn et al., 2000; Gillespie-Lynch et al., 2013; Orr, 2018). Furthermore, research on early communicative behaviors has emphasized gestures as the first means to convey and structure communicative intent. In the context of this reasoning, it is commonly believed that children use gestures several months before they use words.

Some suggest that the emergence of gestures around 9 months offers an explicit means for infants to establish and share reference prior to and during word learning (Volterra et al., 2005; Iverson & Wozniak, 2016). Indexical pointing is discussed as an especially important gestural behavior, forming the basis for primitive deixis (Tomasello et al., 2007; Cochet & Vauclair, 2010; Cameron-Faulkner et al., 2015). Pointing is often cited as a prerequisite skill that leads infants down the path toward linguistic communication (Iverson et al., 1994; Colonnesi et al., 2010). From this perspective, caregivers have the opportunity to label objects of shared interest as infants begin to use gestures (i.e., pointing) to designate objects in the environment, thus, providing a foundation for acquiring new words (Wu & Gros-Louis, 2015).

In accounts of the origin of language, some theorists have supported the idea that language evolved from a primarily gestural mode in the distant past (Hewes, 1973; Arbib et al., 2008; Corballis, 2010; Tomasello, 2010). The study of great apes in captivity has revealed that their gestures are both deliberate and voluntary with distinctive functions that can be adjusted to different contexts (Byrne et al., 2017). In a recent study examining the role of gesture in symbolic development in human and ape infants, researchers documented that gestural symbols in human infants became less frequent than vocal symbols with age, but age-matched ape infants continued to use gestural symbols more frequently across development (Gillespie-Lynch et al., 2013). Interestingly, they found few deictic gestures (that is, pointing gestures) in the apes.

Many prominent researchers in the area of gesture in the great apes have reported that gesture is far more flexible than vocalization in their communication systems, with captive apes communicating primarily through gesture (Tomasello & Zuberbühler, 2002; Pika et al., 2005; Pollick & De Waal, 2007). A key differentiation is that, unlike humans, our ape relatives do not normally assemble their vocalizations into more complex utterances composed of syllables, words, and sentences as humans do (Riede et al., 2005), with perhaps rare and limited exceptions (see e.g., Clay & Zuberbühler, 2009). This viewpoint is supported by findings on relatively successful sign language learning in great apes raised by humans, but almost total failure to learn spoken language in the same circumstances (Gardner & Gardner, 1969; Patterson & Cohn, 1990; Bonvillian & Patterson, 1999; Rivas, 2005; Tomasello, 2007). The claim of greater gestural flexibility in apes has been challenged in recent years, with research suggesting that nonhuman primates may display more flexibility in their vocal production than previously believed (Clay et al., 2015; Cheney & Seyfarth, 2018). The findings suggest that despite the widely acknowledged vocal limitations of the apes, they are capable of modifying both gesture and call usage for different social settings.

### 1.2. Vocalization as a Foundational Activity in Language

Infant non-cry, speech-like vocalizations, or “protophones” are often viewed as one of the primary means by which infants communicate with others. Babies begin vocalizing soon after birth producing an array of protophones. By six months of age, these sounds come to include well-formed syllabic sounds, canonical syllables (Oller, 1980; Locke, 1993). Early longitudinal investigations of newborns indicate that infants can produce spontaneous vocalizations from the first weeks of life (Stark, 1980; Koopsmans-van Beinum & Van der Stelt, 1986; Oller, 2000), with a tendency to produce protophones at very high rates (Nathani et al., 2006). A recent work showed that protophones occur at a much higher rate than cries in both preterm and full-term infants from as soon as infants could breathe on their own (Oller et al., 2019). Although spoken words may not develop until the end of the first year, infants use vocalizations to communicate needs and states of being from the first 3 months of life (Soltis, 2004; Jhang & Oller, 2017). These early spontaneous vocalizations can be viewed as foundational for the development of more complex skills such as canonical babbling and first words. Thus, later speech development is viewed as being built upon the foundation of early vocal production.

The endogenous nature of infant vocalizations—i.e., intrinsically motivated sounds produced for the infant’s own purposes—has gained attention in accounts of language origins. Long et al. (2020) found that infants in the first year produced the majority of vocalizations independent of social engagement, specifically, three times as many endogenous vocalizations as socially-directed ones. Modeling of infant vocal development with computer and robotic simulations has also added support for intrinsic motivation for vocalizing in infancy (Oudeyer, 2005; Oudeyer & Kaplan, 2006). One study emphasized first vocalizations as the result of vocal exploration, i.e., vocal curiosity, which in the modeling led to a progressive mastery of capabilities required for speech, such as phonation (Moulin-Frier et al., 2014).

Despite the findings emerging from infant studies, theories supporting vocalization as foundational for the origin of language have largely focused on the behaviors of nonhuman primates. These investigations have sought to reveal continuities in behavior between human speech and ape vocal communication. There is evidence documenting some, though limited vocal invention and novel vocal learning in great apes (Wich et al., 2009; Lameira et al., 2016). In a study of kiss-squeaks recorded from wild orangutans, researchers found three different variations of the call, suggesting some volitional control (Hardus et al., 2009). Some degree of functional and novel call usage has also been documented in captive chimpanzees (Hopkins et al., 2007).

### 1.3. An Evolutionary-Developmental Perspective

A considerable body of evidence has described the development of communicative behaviors in infancy. We follow a line of thinking from an evolutionary-developmental (evo-devo) framework which emphasizes the widespread tendency for new structural features or capabilities to evolve by modification of developmental patterns (Müller & Newman, 2003;

Bertossa, 2011). Conservation of foundational structures is expected, and natural selection is expected to build upon the foundational structures, keeping developmental sequences consistent with the order of evolved structures and capabilities. In other words, the order of appearance of major structures or behavioral capabilities in development is expected to emerge following evolutionary orders (Newman, 2016; Carroll, 2005). In accord with this line of thought, it seems reasonable to predict that if gesture does form the primary foundation of language, then gestural communication should predominate in early vocal communication in humans.

### 1.4. Specific Aims

The empirical goal of this research is to determine the relative extent to which infants produce communicative or potentially communicative gestures and protophones across three ages in the first year. To our knowledge no prior research has directly compared the rates of gestures and protophones in early infancy. Furthermore, we do not know of any research demonstrating a systematic or clear criterion for the purposes of identifying the potential communicative roles of individual bodily movements. Consequently, it has not been possible to derive very meaningful comparisons across gestural and vocal development. Instead, the study of early infant manual action has yielded limited descriptive accounts for infants prior to six months of age (Fogel & Hannan, 1985), lacking a clear basis for comparison with vocal action and communicative functions.

There is not even a currently widely accepted framework for describing gesture in the first year. Thus, we have known little about how to judge gesture in the first months of life. This gap is especially notable considering well-established procedures for judging both the structure and communicativeness of protophones from birth and across the whole first year. This study is the first to our knowledge to describe a scheme for gestural coding designed to supply a basis for comparison across gesture and vocalization across the first year. Foremost, it provides an opportunity to quantify gesture and vocalization rates for comparison across the first year of life. In doing so, we hope to provide new perspectives on the relative roles of gesture and vocalization in the origin of language and more specifically, their roles in modern human communicative development. Our primary research questions concern the rate of occurrence of potentially communicative gestures compared to the rate of protophones during the first year of life and the relative rates of social directivity of gestures and protophones across the same period.

### 1.5. Hypotheses

1. In accord with the widespread assumption that language is based on gesture, gestural communication should occur at a higher rate than vocal communication in early human development.
2. Again, in accord with the widespread assumption, gestures, more often than protophones, should show a clear sign of constituting intentional communication; in particular, gestures more often than protophones should be accompanied by gaze directed toward another person.

## 2. Methods

### 2.1. Selection of Participants

Data were acquired from archived longitudinal audio-video recordings from the Origin of Language Laboratory (OLL) at the University of Memphis. Approval for the longitudinal research that produced data for this study was obtained from the University of Memphis Institutional Review Board for the Protection of Human Subjects (IRB). All of the participants resided in or around Memphis, Tennessee, and recruitment for this archival data was conducted in child-birth education classes and by word of mouth. All of the infants’ parents completed a written consent form approved by the IRB prior to any of the recordings utilized in the present study. Because parents were recruited during pregnancy, inclusion criteria for participation was initially determined as a normal pregnancy up to the point of recruitment without any detected complications. Typical development (i.e., lack of hearing, vision, language, or other developmental disorders) was confirmed throughout participation in the longitudinal study via parent report during laboratory visits using information such as passed hearing screenings and mastery of developmental milestones at approximately expected ages

The available archived recording sessions included a total of 21 parent-infant dyads from two waves of longitudinal study (see e.g., Oller et al., 2013; Oller et al., 2019). We selected 10 parent-infant pairs (5 male, 5 female infants) balanced for age, gender, recording session type, and recording length for analysis in the present study. Each of the longitudinal recordings included three sessions, each approximately 20-minutes in length, usually yielding a continuous ~60-minute recording. For the current work, we analyzed sessions where parents were instructed to engage their infants in a face-to-face interaction (further discussed as “interactive” sessions) at approximately 4, 7, and 11 months. In the interactive sessions, parents were instructed to interact with their infants vocally as they normally would, whereas the other two recording circumstances during the ~60 minutes required the infant to either be alone (parent present but reading) or be playing separately while a parent was being interviewed by another adult.

We selected infants from the archives with recordings that fit the age criteria to the greatest extent possible with existing vocalization coding from prior work. All but one of the infants were White (Infant 4). All of the infants were learning English as their native language except for Infant 2, who was exposed to German, Spanish, and English at the time of the recordings. Demographics and recording ages for each infant at each session are provided in Table 1. The selection was unbiased with regard to the selected sessions’ being more gestural or vocal activity.

**Table 1.**
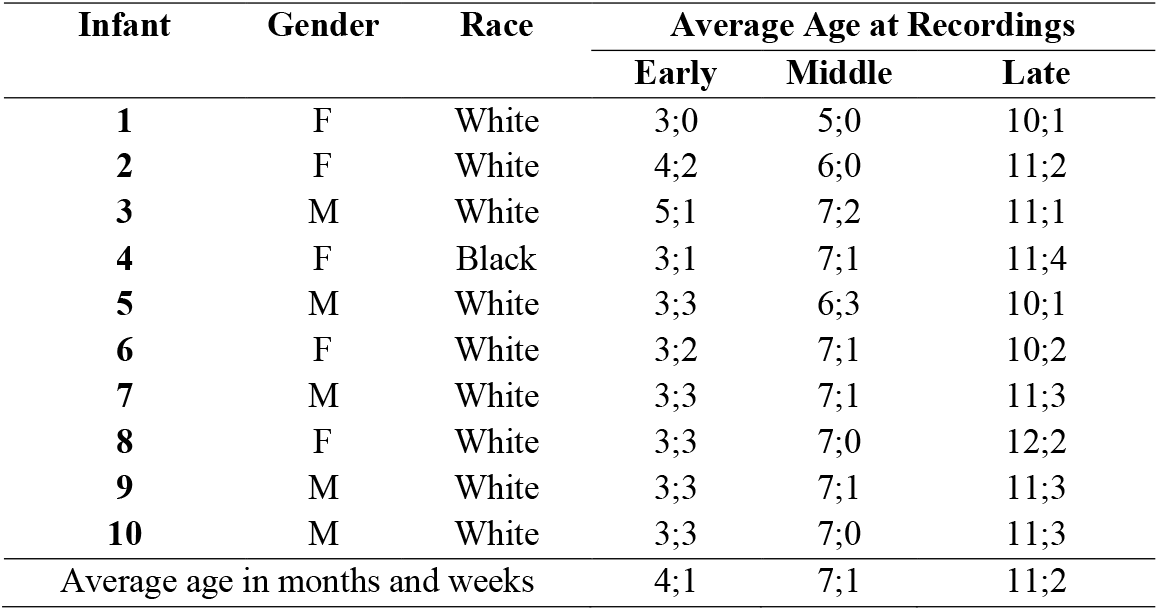
Infant Demographics and Average Ages at Recordings

### 2.2. Procedure for Data Collection

One recording was selected for each infant (total: 10) at each age (total: 3), for a total of 30 recordings. The average length of sessions used for this study was 19 minutes (range: 16 - 20 minutes). All parent-infant dyads were recorded in a sound-treated playroom with toys and books. Both infants and parents wore high fidelity wireless microphones. The playroom was equipped with cameras in each of the four corners, either four cameras or eight (one high camera and one low camera in each corner). The differing number of cameras occurred due to two phases of recording in two different laboratories. The laboratory staff operated the cameras for zoom and tilt from an adjacent control room. Two of the four or eight channels were selected to record the interaction at any given moment. The selection and zooming was intended to afford close up views of the infant’s face, torso, and actions on one of the selected channels and for a broader view of the interaction (including parent) on the other selected channel. More detailed information regarding the laboratory equipment and procedures of recording can be found in previous work from this laboratory (Buder et al., 2008; Oller et al., 2013). Importantly, the instructions to parents for the interactive segments emphasized playing with and interacting with infants in a natural way, allowing for vocal, gestural, and tactile interaction at any time.

### 2.3. Coding Software and Plan

Coding for the current study was completed in AACT (Action Analysis Coding and Training software, Delgado, 2010) which facilitates simultaneous coding of digital video and audio. The AACT program includes two channels of video synchronized with the audio for frame-level accuracy which are synchronized and coordinated with audio represented on a spectrogram-waveform display in TF32 (Milenkovic, 2010). This system permits coding in multiple fields either separately or at the same time. The fields of interest for the present research were protophones (vocal types), gestures, and gaze direction. Aside from gesture coding, data collection was completed in a way that was identical to previous infant vocalization studies conducted in the OLL (Oller et al., 2013; Long et al., 2019; Long et al., 2020; Jhang & Oller 2017). AACT includes a main menu panel for code selection and coding panels which display time boundaries for selected codes (in this case panels for gestural acts, gestural functions, and gaze direction). During coding, the coder selects cursor placements on the TF32 screen and is able to play that selected sequence as many times as desired in a loop. The coder can also drag the cursor on the audio display with corresponding frame accurate changes in the video from both cameras to enable the selection of onset and offset points for gestural events. Also, the coder can move the video screen and audio cursors one frame at a time in either direction with keyboard controls.

[Image removed per BioRxiv preprint policies]

**Figure 1.** Example of AACT coding program in action. This figure illustrates two video displays synchronized and coordinated with an audio display including both waveform and spectrogram, as seen on the left screen. The right screen shows the menu panel on the left and three overlapping coding panels (in this case panels for gestural acts, gestural functions, and gaze direction) on the right. The coder is able to select cursor placements on the audio screen and then play that selected sequence as many times as desired in a loop. The coder is also able to drag the cursor on the audio display with corresponding frame accurate changes in the video from both cameras to enable the selection of onset and offset points for gestural events. Also, the coder can move the video screen and audio cursors one frame at a time either direction with keyboard controls.

### 2.4. Vocalization Coding

Data for vocalizations had been coded in prior research (e.g., Oller et al. 2013, Oller et al. 2019), with nearly identical procedures of coding having been utilized for all infants. The primary focus of this previous work was speech-like vocalizations, focusing on phonatory properties of sounds. This approach resulted in three primary types being considered: vowel-like, growl-like, or squeal-like sounds. Oller (2000) refers to these types of vocalizations as protophones (including both canonical and pre-canonical sounds), the presumed precursors to speech. Cries, laughs, and whimpers were also coded but not included for analysis in the current study. Coders were instructed to use repeat-observation to assign boundaries in TF32 at the onset and offset of each protophone. The cursor placements at coding of each utterance in TF32 were recorded in AACT, specifying the duration of each speech-like sound. The boundaries were assigned using a “breath-group” criterion (Lynch et al., 1995); an utterance begins with phonation during exhalation (i.e., egress) and ends with the termination of phonation, often accompanied by inhalation (i.e., ingress). In accord with this criterion, a new utterance can begin after the observed breathing pause. After the coder determines where the onset and offset of each utterance occurs, a protophone type is selected from the list of options on the coding panel, and in a subsequent pass, gaze direction is assigned (directed to a person or not). Extensive details on the vocalization and gaze coding and the associated intercoder agreement can be found in prior publications, especially in the accompanying supplementary documents (e.g., Oller et al. 2013; Jhang et al., 2017).

### 2.5. Gesture Coding

#### 2.5.1. Global Categories for Gesture Coding

We developed a coding scheme to categorize acts during the first year of life that could be considered precursors to signs (that is, precursors to gestural symbols or gestural performatives). The gesture coding labels are organized in terms of our proposed hierarchy of actions that form a foundation for language-like gestures. We designate four global categories where each successive category is interpreted as more communicatively advanced and more language-like than the prior ones. These include: 1) Utilitarian Acts, 2) Non-social Gestures, 3) Universal Social Gestures, and 4) Conventional Gestures. We define *Utilitarian* Acts as those actions that are simply world-exploratory or manipulative, without any inherently social communicative function. For example, if a child reaches for and obtains a toy, thus serving his or her non-communicative goal, the event is coded as a Utilitarian Act. Actions included in this category were coded in the initial real-time coding pass, and this coding provided a basis for locating utterances in subsequent passes, but the real-time codes were excluded from the analyses below.

The remaining three global gestural categories are intended to include all the actions that could conceivably be interpreted as communicative or those that could be expected to be brought into the service of communication at some later point in development. *Non-social Gestures* are gestural acts that are not merely utilitarian and are also not inherently communicative, although they have *the potential for being utilized communicatively.* For instance, rhythmic hand banging and foot tapping are examples of Non-social Gestures. *Non-social Gestures* are akin to babbling in the vocal domain, actions that can eventually be brought to the service of communication, but that are not yet intentionally communicative. *Universal Social Gestures* include any act with an inherently social communicative intent but with no indication of having been learned from specific cultural experience; for example, an extended flat hand to indicate refusal is a Universal Social Gesture. Also, reaching upward with both hands when an infant wishes to be picked up can be thought of as a Universal Social Gesture. *Conventional Gestures* are those that are presumed to be culturally specific or culturally transmitted—acts that have a discernible communicative function, such as a hand wave to convey “hello” or “bye-bye”, clapping in celebration, or thumbs up to indicate approval or agreement. In the vocal domain, these categories facilitate comparison across vocalization and gesture, since the vocalization coding also presumes a hierarchy of complexity where each successive level is more communicatively advanced than the earlier one. Non-social acts are least communicative and least learned, and Conventional acts are most communicative and most learned. Protophones and speech can similarly be subcategorized as Non-social (including both pre-canonical and canonical babbling) and Conventional (speech). Universal social acts fall between Non-social and Conventional levels in the degree to which they are language-like because they are communicative but (for example) require no associative learning (presumably only motoric learning is required) and since they are (we presume) recognizable and interpretable as to intended function to potentially everyone around the world. Universal Social Gestures are capable of transmitting certain critically important communicative functions such as refusal, request, and designation (pointing). In the exclusively vocal domain, these functions seem impossible unless symbols (words) are involved, and even then vocal transmission is more complicated than gestural. Consider designation, which is possible with words (“look to your left and notice an orange object”), but not with protophones. Similarly, other functions that can be transmitted with gesture universally (refusal or request for example) cannot be uniquely transmitted in vocalization without symbolism, although prosodic features of vocalization and/or facial affect can emphasize or modulate the flavoring of such functions in any modality of transmission—when they are transmitted by gesture or in signed or vocal symbols. There is thus a gap in the potential for communication transmitted with protophones, a gap that can be at least partially filled by Universal Social Gestures, which consequently provide a scaffold for early communication, supporting and coordinating such gestures with vocalization and facial affect as language begins to emerge. This special capability of Universal Social Gestures may account for the widespread opinion that human language is founded in gesture.

#### 2.5.2. Primary Categories for Gesture Coding

The primary coding for the current research was conducted by the first author. Based on initial attempts at creating a workable coding scheme for gesture, with reasonable inter-coder agreement, we determined a clear distinction needed to be drawn between gestural *acts* and the potential *intent*, or communicative function, associated with each action. This distinction is also required between vocal acts and their communicative functions. For example, reaching in itself describes an action, but it does not characterize the communicative function of the act. An infant could reach with an extended arm to show an object, to request an object, to try to get the attention of another person, or to offer or accept something. Our coding scheme included two fields, allowing judgements for the individual gestural act in one case and for the communicative function of each act in the other.

The gestural acts and gestural functions were coded for each ~19-minute segment in three separate passes using both audio and video. A first pass using real-time observation was completed in AACT to allow the coder to acquire an overview of the recorded segment and its interactions and to mark the approximate location of infant gestures, labeled by way of key strokes, each corresponding to one of the following global codes: Utilitarian, Non-social, Universal Social, or Conventional. In the second pass, the primary coder used repeat-observation to designate boundaries for individual gestural acts in the TF32 module (the acoustic display) in AACT at the onset and offset of each action, specifically designating the point of the movement beginning the act through the point where the event reached its full extension. For example, the onset of an index-finger point begins when the arm, hand, or finger moves into motion from rest until the point where it reaches its full extension. In cases of rhythmic movements such as hand banging, boundaries are determined using a criterion similar to the breath-group criterion for vocalizations. Thus, duration of any cluster of actions (such as the individual strokes of rhythmic hand banging), deemed to constitute a gesture that occurs before an observed gestural pause (treated similarly to pauses in speech), specifies the duration of the gesture. The cursors in TF32 were adjusted to make each bounding decision, which sometimes required that the cursors first be set as much as 1000 ms before the onset and after the offset for extended viewing of the event using playback features in AACT. Then the cursors could be moved (by dragging the cursor in the acoustic display with corresponding frame accurate changes in the video or by keystrokes that could move both audio and video displays one frame at a time) in to the actual boundaries of the gesture specifying the onset and offset points. A keystroke indicating a particular gestural act would then be recorded in AACT to indicate both onset and offset times for that gesture.

Once the onset and offset boundaries had been determined, we used the bounded time frames that had been assigned to the gestural acts to automatically create placeholders in a new coding panel based on a special AACT menu feature. The placeholders were recorded sequentially in a new coding panel and allowed the coder to categorize all the bounded events in a third pass, where a gestural function was designated for each gestural act. The second and third coding passes for gestural acts and gestural functions used the detailed category labels as listed in Table 2 and Table 3, respectively.

**Table 2.**
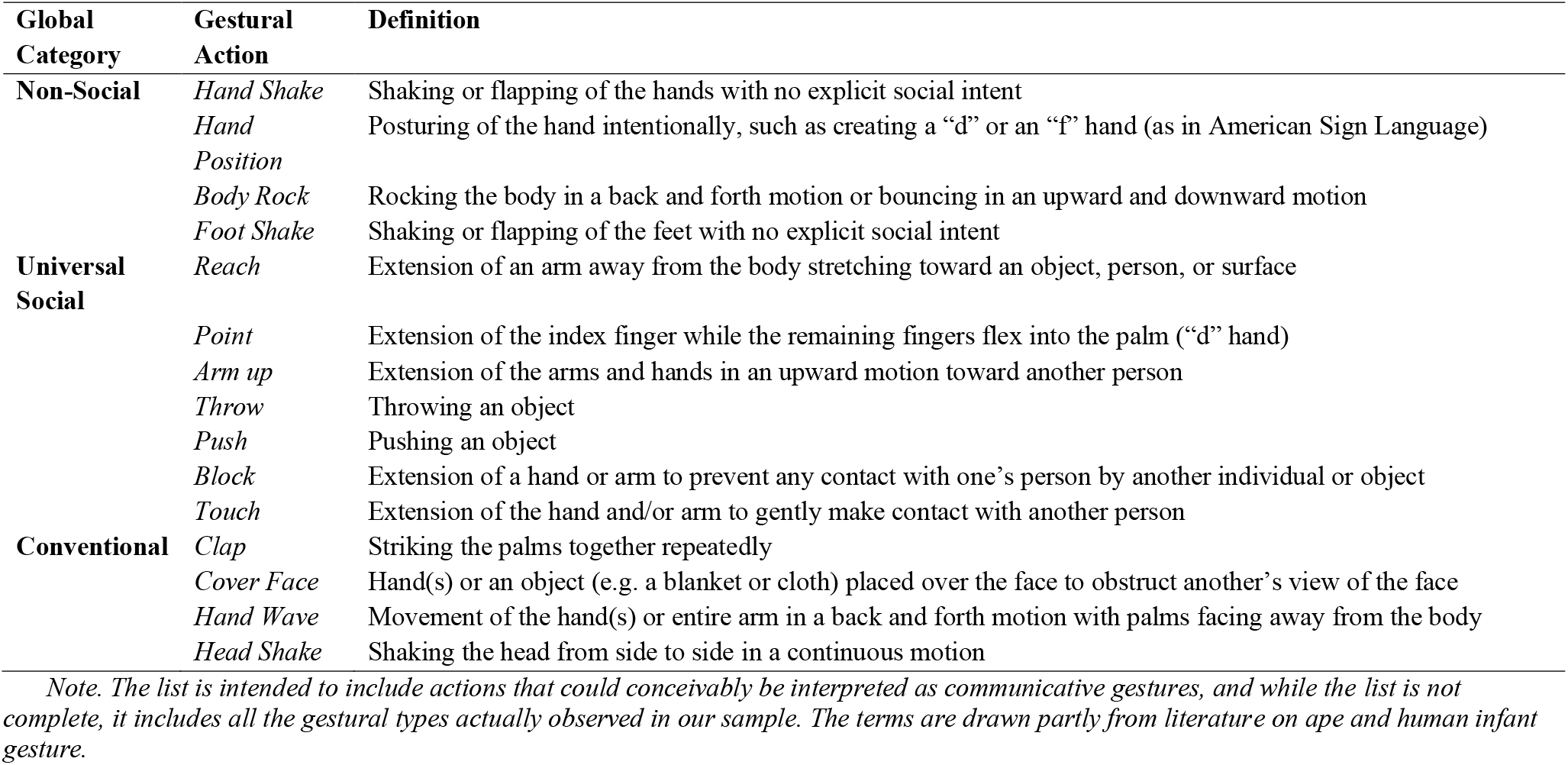
Gestural Action Categories Used in the Present Study

**Table 3.**
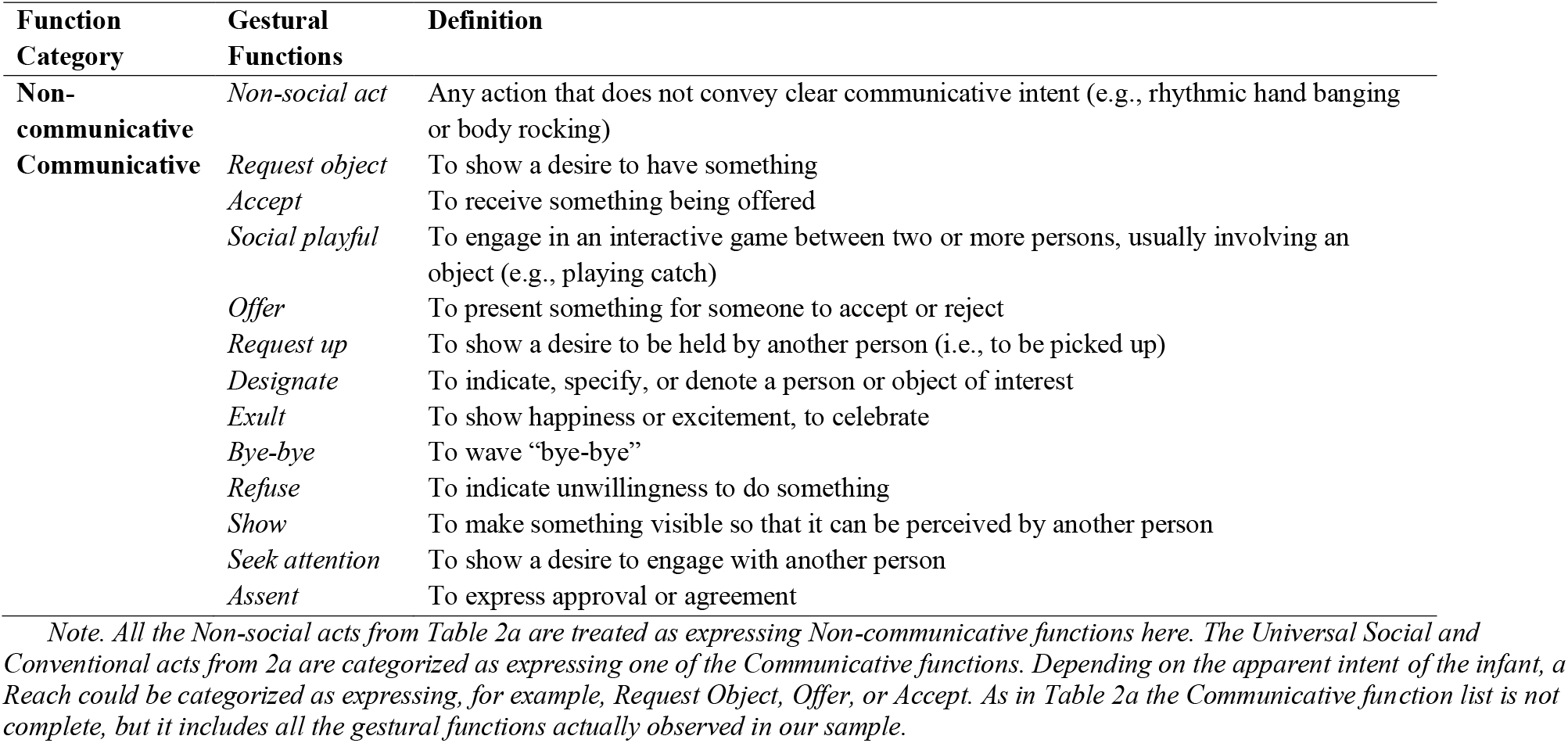
Gestural Function Categories Used in the Present Study

### 2.6. Gaze Coding

In a fourth pass, we coded the gaze direction of infants during each vocalization and during each gesture where at least one of the two video views allowed such judgement. We coded whether infants produced a protophone or a gesture while looking at another person (i.e., a socially-directed communication) or while not looking at another person (i.e., a non-socially-directed communication). The primary coder used placeholder time frames for both protophones and gestures to code for gaze direction during those time frames. Of course, the gaze direction for protophones had been previously coded during other studies (e.g., Oller et al. 2013) and those judgements were used in the current study. Following the same coding procedure as in the third pass of coding gesture, coders clicked on each individual event (i.e., coded gesture or vocalization) and then coded the direction of the infant’s gaze during the bounded action. The coders adjusted the cursors so that playback for coding started 50 ms before the event onset and continued through 50 ms after the offset. This procedure was designed to ensure the entire event (all its corresponding video frames) would be viewed before coding the directedness of the infants’ gaze. The categories available for coding were 1) directed toward another person, 2) not directed toward another person, or 3) can’t see (sometimes the infant’s eyes were not visible in either camera view).

In all cases where coders used the placeholder feature to code in a new field (for example, to code in the gaze direction field based on placeholders based on coding from the gesture field), they were blind as to the codes that had been designated in the prior field. Each field was thus coded without information regarding any of the prior coding in other fields.

### 2.7. Coder Training

Two graduate students in Communication Sciences and Disorders at the University of Memphis were trained for real-time and repeat-observation coding for gestural acts, gestural functions, and gaze direction. The training began with a lecture by the first author on the gesture coding scheme described above. During the lecture, coders were presented in AACT with real examples of infant gestural acts previously coded by the first author and confirmed by the last author, and all of which either met a consensus standard for one of the gesture categories or displayed ambiguities of possible coding judgements that were deemed instructive for the training. Gaze direction training was similar. Once the training was completed, the coders followed the same coding procedure as the first author using the criteria outlined in the gestural coding scheme and the gaze direction scheme. The first author selected 12 recordings semirandomly from the 30 recordings. A five-minute segment was randomly selected from within each of the 12 recordings.

### 2.8. Coder Agreement

Four samples came from each of the three ages and all ten infants were represented in the agreement samples. Neither of the agreement coders had any knowledge of the hypotheses for this study. In addition, they were blinded to the outcomes of the coding conducted by the first author. The agreement coding revealed high agreement for the total number of gestures identified in the five-minute segments for the first and second agreement coders (*r*= 0.92, *r*= 0.81, *N*= 12) with respect to the primary coder. The agreement coders also showed a high level of agreement (*r*= 0.80) with each other. For agreement on the difference between the number of gestures coded and the number of gestures with gaze directed to a person, the first and second agreement coders showed good agreement (*r*= 0.92, *r*= 0.73) with respect to the primary coder. There was also good agreement between the two agreement (*r*= 0.72) regarding this difference.

Data from all three coders showed at least a doubling of the rate of gesture from the first to the second age, with an increase ranging from approximately 50% - 60%. The coders also showed an increase in the amount of gesture from the first age to the third age, with an increase ranging from 22% - 65%. Across the entire set of five-minute segments in the agreement samples the three coders showed very similar numbers of person-directed (by gaze) gestural actions (proportion of gestures out of all communicative events both gestural and vocal: 27%, 30% and 30% for the three coders respectively). For all three the coders, 13%– 14% of the gestures were deemed to be directed to a person, whereas the degree of directivity for protophones was > 27%. As will be seen from the Results, the agreement data suggest that if the entire data set had been coded by either of the agreement coders instead of the primary coder, none of the conclusions associated with the results reported below would have changed.

## 3. Results

### 3.1. Distribution of Gestures and Protophones

The data on protophone and gesture rates across all 30 recordings for the three ages of the first year revealed vastly more protophones (3,730) than gestures (738). Figure 2 presents the data for the three ages, showing gestures occurred infrequently (less than one per minute) at the earliest age, increasing to 1.6 – 2 per minute at the later ages. In contrast, protophones occurred more than 8 times per minute at the youngest age, falling to approximately 5 per minute at the oldest age.

**Figure 2.**
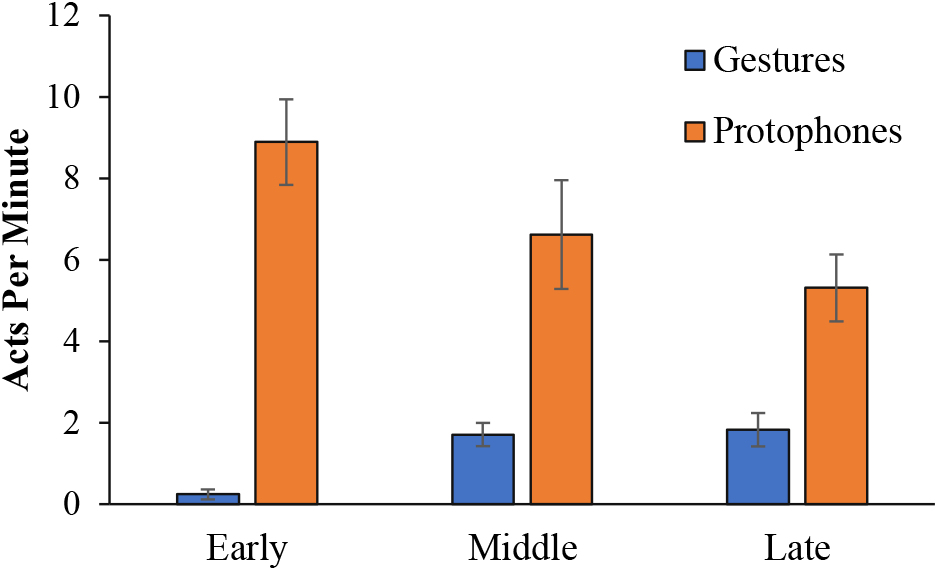
Gesture and protophones per minute across the first year. This figure shows protophones and gestures per minute across the three ages in our sample (error bars show standard errors).

To test the number of protophones vs. gestures at each age, we used paired *t*-tests with a Bonferroni adjusted alpha level of .017 (.05/3). Paired *t*-tests revealed significant differences in the number of infant gestures and protophones across all three ages. Means showed more protophones (*M*= 154.8, *SD*= 67.58) than gestures (*M*= 4.8, SD = 7.31) at the early age, *t*(9) = 7.44, *p*<.001, *d*= 3.12, 95% CI [1.82, 4.43], more protophones (*M*= 122.7, *SD*= 76.29) than gestures (*M*= 33.8, *SD*= 18.48) at the middle age, *t*(9) = 3.40, *p*=.008, *d*= 1.60, 95% CI [0.59, 2.61], and more protophones (*M*= 95.5, *SD*= 44.9) than gestures (*M*= 35.2, *SD*= 23.54) at the late age, *t*(9) = 4.30,*p*<.01, *d*= 1.69, 95% CI [0.66, 2.70]. All findings were statistically significant, and the effects were all large. The results strongly contradict the widespread expectation that gesture should occur earlier than protophones.

To test the effects of Age, we used multiple paired *t*-tests with a Bonferroni adjusted alpha level of .008 (.05/6). The tests indicated a significant increase in the number of gestures from the earliest age to the middle age (*M*= 29, *SD*= 18.19); *t*(9) = 5.04,*p*<.001, *d*= 1.51, 95% CI [0.56, 2.47], and from the earliest age to the latest age (*M*= 30, *SD*= 20.45); *t*(9) = 4.70, *p*<.001, *d*= 1.67, 95% CI [0.71, 2.64]. There was no significant change in the number of gestures from the middle age to the late age, *t*(9) =.159, *p*=.877. Our results (see Figure 2) revealed a decrease in protophones per minute from the earliest to latest age in our sample. In spite of the apparent trend, these differences were not statistically significant: early to middle ages, *t*(9) = 1.19, *p*=.266, middle to late ages, *t*(9) = 1.44, *p*=.185, and early to late ages, *t*(9) = 2.50, *p*=.034. This trend will be taken up in the Discussion.

The great majority of the events observed in our sample were not directed toward another person in *either* modality. Figure 3 shows, however, that the rates of gaze direction were greater for protophones than gestures at each of the three ages. To test the degree of directivity for protophones and gestures at each age, we used paired *t*-tests with a Bonferroni adjusted alpha level of .017 (.05/3). The dependent variable for these analyses was the number of acts in either modality that were directed to a person as a proportion of the total number of acts in both modalities. At the earliest age, protophones (*M*=.43, *SD*=.18) were more often directed than gestures (*M*=.003, *SD*=.01; *t*(9) = 7.43, *p*<.0001, *d*= 1.41, 95% CI [.43, 2.39]; for the middle age, protophones (*M*=.28, *SD*=.15) were more often directed than gestures (*M*= ..05, *SD*=.08; *t*(9) = 5.97, *p*<.0001, *d*= 1.13, 95% CI [.18, 2.07]; and for the latest age, protophones (*M*=.21, *SD*=.17) were more often directed than gestures (*M*=.04, *SD*=.03; *t*(9) = 3.17, *p*=.011, *d*= 1.04, 95% CI [.11, 1.97]. All tests were statistically significant after Bonferroni correction, and all the effects sizes were large.

**Figure 3.**
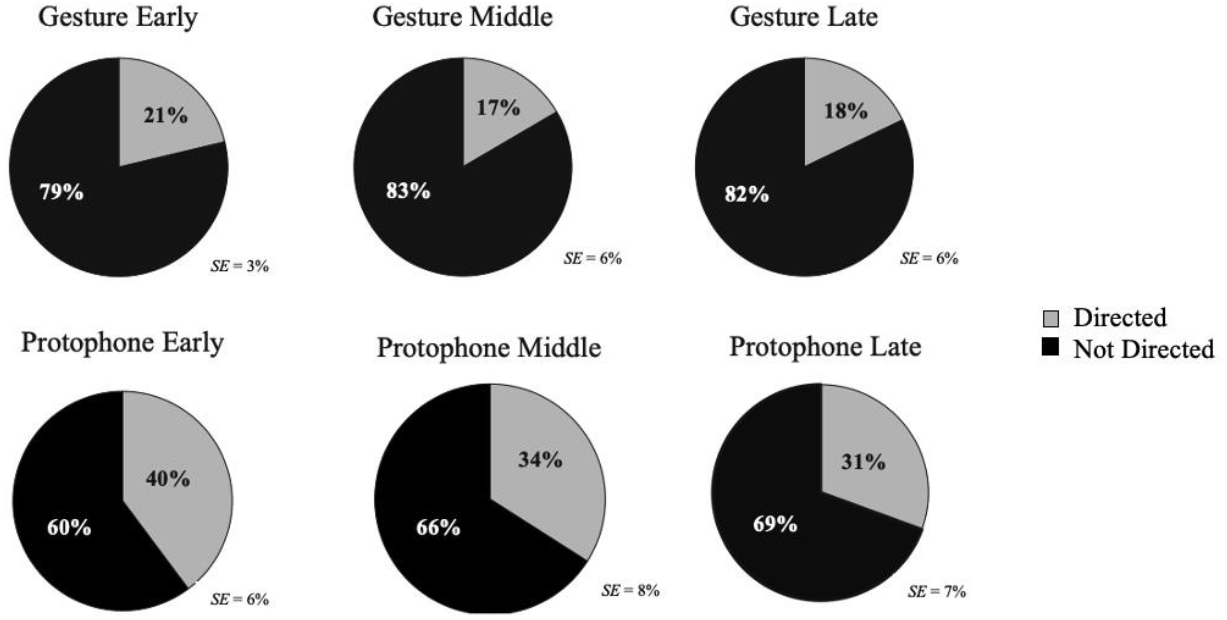
Directivity of gestures and protophones at each age. This figure shows percentages of gestures and protophones that were directed and not directed by gaze at each of the three ages.

### 3.2. Distribution of Gesture and Protophone Types

Table 4 shows the distribution of the three hierarchical categories for gesture. There were roughly the same number of Non-Social and Universal Social Gestures observed at the early and middle ages respectively. Furthermore, Non-Social and Universal Social Gestures were by far the most frequent gesture types at the early and middle ages. At the late age, Universal Social Gestures predominated, accounting for ~ 60% of all gestural events. One might have imagined that pointing (a clearly Universal Social Gesture) would have accounted for much of this difference, but it did not. Pointing only occurred a grand total of 12 times in the 600 minutes of coded recording, and these events were mostly from one infant at 11 months. Both at the middle and late ages, the great bulk of the cases of Universal Social Gestures involved reaching as if to request an object (not actually reached by the infant), offering an object to another person by reaching, or accepting an object by reaching.

**Table 4.** Distribution of Gesture Types

| Gesture Type | Infant Age Group |  |  | Total |
| --- | --- | --- | --- | --- |
|  | Early | Middle | Late |  |
| **Non-Social** | 25 | 162 | 117 | 304 |
| **Universal Social** | 23 | 161 | 211 | 395 |
| **Conventional** | 0 | 16 | 24 | 40 |
| **Total** | 48 | 339 | 352 | 739* |

*Note.* \*This number represents the total number of gestures observed across the 600 minutes of recording.

Conventional Gestures, the types that seem most language-like because they have to be learned, consisted of a total of 40 events, 24 of which occurred at the late age. The tokens of Conventional Gestures that actually occurred at 11 months included only four types: no-no (head or finger shaking), hand clapping, bye-bye (hand waving), and face covering. All of the Conventional Gestures documented at the middle age were associated with a peekaboo game with one infant only, where we counted as Conventional Gestures the infant covering her face and later uncovering it when the father said “peekaboo”. One might question whether any peekaboo game could exist without covering the eyes and uncovering them, and consequently one might question whether these gestures belong in the Universal Social category rather than the Conventional category. But we know of no empirical evidence that supports any claims suggesting peekaboo is a phenomenon that emerges naturally across all cultures, or that it occurs as an interactive game across all cultural contexts. Still, it does occur across many cultures (e.g., Fernald & O’Neill, 1993). We operate under the assumption that infants learn this behavior through caregiver-child play.

One might have predicted, based on the literature’s focus on gesture in the development of language, that there would be extensive utilization of Conventional Gestures by 11 months, and presumably more Conventional Gestures than Conventional Vocalizations. Although the available coding of protophones did not specify them in terms of the three hierarchical categories (social and non-social usage of vocalizations had not been coded), we deemed it informative for the present study to evaluate the number of tokens of Conventional Vocalizations across the three ages. Our count of tokens of Conventional Vocalizations yielded 50, more than the number of Conventional Gestures that were observed. They included mama, dada, bye-bye, no-no, yum-yum, mmm (tastes good), and yeah. Overall, neither Conventional Gestures nor Conventional Vocalizations were frequent compared to the much larger numbers of Universal Social and Non-social acts.

## 4. Discussion

The present study provides the first direct comparison of the rates of gesture and protophones across the first year. Infants used more than five times as many protophones as gestures. A significant imbalance favoring protophones occurred across all three ages. The vocal predominance applied whether the communication was directed or not directed by gaze toward another person, with the substantial majority of both vocal and gestural activity having non-person-directed gaze—this pattern of predominantly endogenous rather than social acts with potentially communicative effect has been observed previously in the vocal domain (Long et al., 2020). The present data also show that protophones were produced more often with person-directed gaze than gestures. There were 1,320 cases across the first year where a protophone was directed by gaze compared with 129 cases of directed gaze for gesture. Furthermore, protophones were almost twice as likely to be person-directed. If we take gaze direction as an indicator of communicative intent, we can conclude that not only did protophones occur much more frequently than gestures in general, but there were far more events that were intentionally communicative in the vocal domain.

Our research dramatically contradicts the assumption that gesture is predominant in early infant communication. Because infants produced a noticeably greater number of protophones from an early age, it is reasonable to suggest that the roots of language are deeply grounded in the vocal domain, supporting the notion that the vocal ability and inclination to produce these sounds constitute a fundamental first step toward more complex language skills. The results suggest vocalization is a more important foundation for language than gesture.

The great majority of gestures observed were not the ones expected based on the common suggestion that language is founded in gesture. Pointing, which is known to form a foundation for word learning, occurred far less frequently than we expected. Only 12 pointing events were observed (<2% of observed gestures), with only two of ten infants having pointed during these naturalistic interactions in a laboratory setting designed to resemble a child’s playroom. Reaching (including merely reaching toward an object and not obtaining it without help, offering an object, or accepting an object) accounted for 48% of all gestures, suggesting that the “declarative” function of pointing, thought to be so important as a foundation for language, through 11 months of age, occurs far less frequently than the instrumental functions associated with reaching. Given its obvious importance, we were surprised at the infrequent occurrence of pointing. Only one other category of gesture occurred fairly frequently. Rhythmic hand shaking, which we interpreted as a Non-social Gestural act similar to reduplicated babbling, accounted for 35% of all gestures and even accounted for 26% at 11 months. Together, hand shaking and reaching accounted for 83% of gestures.

Both Conventional Gestures and Conventional Vocalizations were infrequent in the first year. Forty cases of Conventional Gestures were observed along with 50 instances of spoken words. Both the Conventional Gestures and the words were overwhelmingly performatives, that is, they constituted illocutionary acts such as greeting (waving, saying hello), celebrating (clapping, saying hooray), or refusing (head shaking, or saying no), rather than semantic acts of reference, such as naming an object or describing an event. Thus, the Conventional acts in both modalities were overwhelmingly dyadic, constituting communications between two parties with respect to each other, rather than being triadic communications where the two parties jointly referred to a third entity through a conventional symbolic act.

The relative tendency to communicate in the vocal domain compared to the gestural domain in early life is not only important in building our understanding of the emergence of the speech capacity in modern human development, but it also offers insights into the likelihood that vocal communication predominated in the evolution of language. We reason that if language indeed originated from gestural use, gestural activity should have occurred to a far greater extent than we saw in this study.

In creating and using a theoretically informed scheme for coding gestural acts and functions, we found that there were roughly the same number of Non-Social and Universal Social Gestures at the early and middle ages. At the late age, Universal Social Gestures predominated and constituted 60% of all gestures, highlighting an advantage of the gestural modality. The advantage is that Universal Social Gestures communicate a clear intent, often of a deictic sort, indicating something the infant wants or something the infant wants another person to look at. On the other hand, vocalizations must first become symbolic in order to serve deictic functions. For example, an infant can gesture by extending his or her arm(s) to signal the desire to be picked up, but there is no equivalent in the vocal domain without words (“pick me up” or just “up”).

Of course, vocalization *can* transmit emotional valence, and so can be used universally to modulate the affective tone of communication, although words (or gestures) are required to designate the entity that is the focus of the affect. It is universal for caregivers to recognize cry and fussy sounds (each protophone type can be flavored by intonation or other acoustic modulations to convey affect), but that universality does not designate what the fussing is about. Approximately 15% of protophones in laboratory recordings labeled as fussy can be judged as such from sound alone (Jhang & Oller, 2017), and so it can be concluded that vocalization does commonly play a role in transmitting affect even in infancy. Thus, protophone prosody can assist in flavoring the affect of communication when it accompanies Universal Gestures, but it cannot supplant the gestures, because it cannot transmit the deictic information that is so natural to the gestural domain. Facial affect can also be utilized to modulate the emotional tone of communication in either gesture or vocalization, in a way that is similar to modulation of emotional tone by the prosody of vocalization.

Another way vocalization can be used to assist in gestural communication is in supporting attention seeking. Infant gestural acts such as pointing or reaching are often accompanied by vocalizations to attract the attention of the listener. But even in these cases, the vocalizations (unless they are words) cannot serve the deictic functions that are natural to the gestural domain.

An important feature of protophones is their functional flexibility (Oller et al. 2013, This feature contrasts sharply with the functions associated with Universal Social Gestures. The latter involve identifiable gestural types each of which is associated naturally with a particular function or class of functions. Pointing, for example, is naturally associated with designation. In contrast, no protophone type has a universal function—all protophones can be produced to serve multiple functions. It is a critical feature of protophones to be free to serve functions with all possible valences, because if they were not free in that way, they could not form a foundation for words, which are by definition free of any particular illocutionary function. Every protophone, just as every word, must be free to express any emotional state, any illocutionary function, or importantly, must be free to be produced for no purpose other than interest in the sound itself. The Universal Social Gestures are unique precisely because they do not possess this flexibility. Their advantage is that they are consequently capable of serving a particular function or set of functions with great consistency.

Of course, both gesture and vocalization are capable of developing into full-fledged language with full functional flexibility. Both modalities include actions that are not inherently tied to particular universal functions. Non-Social Gestures are in this sense like vocal babbling. In both cases (Non-Social Gestures and protophones) the infant explores actions that are free of particular function and can thus be adapted at a later point by learning to form Conventional acts.

The flexibility of protophones is emphasized by recent results from coding of laboratory recordings, suggesting ~75% of infant protophones are produced without social directivity (Long et al., 2020). This fact suggests protophone production is largely endogenous. It has been reasoned that protophone production offers caregivers information about an infant’s well-being; i.e., a caregiver may interpret the sound of an infant vocalizing (whether in interaction or alone) as a signal of their wellness, even if the caregivers are not fully attending. The same kind of reasoning may be thought to apply to gestural babbling, i.e., to Non-Social Gestures.

Infants do not need to have anyone looking at them for vocal communication, although it may help. In contrast, gesture is inherently visual and requires the visual attention of the receiver, even if just with a glance, in order for communication to be successful. Yet, when they gestured, babies at all three ages rarely looked to see if the potential receiver was looking at them. The fact that directivity appeared to be at its highest (though not statistically significantly) at the earliest age may be an artifact. Specifically, the parents at the earliest age were often directly in the infant’s face during the interactions. The youngest infants were not yet crawling or walking and thus not able to move away from parents.

For the vocal domain, the apparent (though not statistically significant) decrease in the rate of vocalization across the first year may not reflect an actual decrease in infant volubility. Previous studies have noted a similarly lower rate of vocalization across age as measured in utterances per minute, but not as measured in syllables per minute (Iyer & Oller, 2008; Iyer et al., 2016). Infants appear to produce longer utterances with more syllables per utterance as they grow older.

## 5. Conclusion

Overall, we found that vocalization is overwhelmingly predominant over gesture in communication of the first year. The vocal predominance applied whether the communication was directed or not directed by gaze toward another person. Our results dramatically contradict the assumption that gesture is predominant in early infant communication, and thus suggest that gesture may not form the primary foundation of language. On the contrary, the results are consistent with the suggestion that vocalization is, and always has, formed a more important foundation for language than gesture.

## 6. Acknowledgements

The research for this manuscript was funded by NIH Grants DC006099 and DC0011027 from the National Institute on Deafness and Other Communication Disorders and by the Plough Foundation, which supports D. Kimbrough Oller’s Chair of Excellence.

## Ethics Statement

This study was conducted in accordance with the recommendations of the Institutional Review Board (IRB) guidelines for the University of Memphis (https://www.memphis.edu/research/researchers/compliance/irb/index.php) with written consent from all parents of infants in the study in accord with the Declaration of Helsinki.

## Author Contributions

**Megan Burkhardt-Reed:** Conceptualization, Methodology, Investigation, Formal analysis Writing-Original draft preparation, Visualization; **Helen Long:** Conceptualization, Methodology, Writing-Reviewing and Editing; **Dale Bowman:** Formal analysis; **Edina Bene:** Project administration; **D. Kimbrough Oller:**Supervision, Writing-Reviewing and Editing.

## References

Armstrong, D. F., & Wilcox, S. E. (2007). The gestural origin of language. Oxford University Press. https://doi.org/10.1093/acprof:oso/9780195163483.001.0001

Bates, E., Camaioni, L., & Volterra, V. (1975). The acquisition of performatives prior to speech. Merrill-Palmer Quarterly, 21(3), 205–226.

Bates, E., Benigni, L. Bretherton, I., Camaioni, L., & Volterra, V. (1979). The emergence of symbols: Cognition and communication in infancy. Academic Press.

Bertossa, R. C. (2011). Morphology and behaviour: Functional links in development and evolution. Philosophical Transactions of the Royal Society B: Biological Sciences, 366(1574), 2056–2068. http://doi.org/10.1098/rstb.2011.0035

Bonvillian, J. D., & Patterson, F. G. (1999). Early sign-language acquisition: Comparisons between children and gorillas. In S.T. Parker, R.W. Mitchell, & H. L. Miles (Eds), The mentalities of gorillas and orangutans: Comparative perspectives (p. 240–264). Cambridge University Press. https://doi.org/10.1017/CBO9780511542305

Buder, E. H., Chorna, L. B., Oller, D. K., & Robinson, R. B. (2008). Vibratory regime classification of infant phonation. Journal of Voice, 22(5), 553–564. http://doi.org/10.1016/i.ivoice.2006.12.009

Byrne, R. W., Cartmill, E., Genty, E., Graham, K. E., Hobaiter, C., & Tanner, J. (2017). Great ape gestures: Intentional communication with a rich set of innate signals. Animal Cognition, 20(4), 755–769. https://doi.org/10.1007/s10071-017-1096-4

Call, J., & Tomasello, M. (Eds.). (2007). The gestural communication of apes and monkeys. Taylor & Francis Group/Lawrence Erlbaum Associates.

Cameron-Faulkner, T., Theakston, A., Lieven, E., & Tomasello, M. (2015). The relationship Between infant holdout and gives, and pointing. Infancy, 20(5), 576–586. https://doi.org/10.1111/infa.12085

Carroll, S. B. (2005). Evolution at two levels: On genes and form. PLoSBiol, 3(7), e245. http://doi.org/10.1371/iournal.pbio.0030245

Cheney, D. L., & Seyfarth, R. M. (2005). Constraints and preadaptations in the earliest stages of language evolution. The Linguistic Review, 22(2-4), 135–159. https://doi.org/10.1515/tlir.2005.22.2-4.135

Cheney, D. L., & Seyfarth, R. M. (2018). Flexible usage and social function in primate vocalizations. Proceedings of the National Academy of Sciences, 115(9), 1974–1979. http://doi.org/10.1073/pnas.1717572115

Clay, Z., & Zuberbühler, K. (2009). Food-associated calling sequences in bonobos. Animal Behaviour, 77, 1387–1396. https://doi.org/10.1016/j.anbehav.2009.02.016

Clay, Z., Archbold, J., & Zuberbühler, K. (2015). Functional flexibility in wild bonobo vocal behaviour. PeerJ, 3, e1124. http://doi.org/10.7717/peerj.1124

Cochet, H., & Vauclair, J. (2010). Pointing gestures produced by toddlers from 15 to 30 months: Different functions, hand shapes and laterality patterns. Infant Behavior and Development, 33(4), 431–441. http://doi.org/10.1016/j.infbeh.2010.04.009

Colonnesi, C., Stams, G. J. J., Koster, I., & Noom, M. J. (2010). The relation between pointing and language development: A meta-analysis. Developmental Review, 30(4), 352–366. http://doi.org/10.1016/J.DR.2010.10.001

Corballis, M. C. (2009). The Evolution of Language. Annals of the New York Academy of Sciences, 1156(1), 19–43. http://doi.org/10.1111/j.1749-6632.2009.04423.x

Corballis, M. C. (2010). The gestural origins of language. Wiley Interdisciplinary Reviews: Cognitive Science, 1(1), 2–7. http://doi.org/10.1002/wcs.2

De Condillac, É. B. (2001). An essay on the origin of human knowledge (H. Aarsleff, Trans.). Cambridge University Press. (Original work published 1756)

Delgado, R.E., Buder, E.H., & Oller, D.K. (2010). AACT (Action Analysis Coding and Training). Intelligent Hearing Systems, Miami, FL.

Fernald, A., & O’Neill, D. K. (1993). Peekaboo across cultures: How mothers and infants play with voices, faces, and expectations. In K. MacDonald (Ed.), SUNY series, children’s play in society. Parent-child play: Descriptions and implications (p. 259–285). State University of New York Press.

Fogel, A., & Hannan, T. E. (1985). Manual actions of nine-to fifteen-week-old human infants during face-to-face interaction with their mothers. Child Development, 1271–1279.

Gardner, R. A., & Gardner, B. T. (1969). Teaching sign language to a chimpanzee. Science, 165(3894), 664–672. http://doi.org/10.1126/science.165.3894.664

Gillespie-Lynch, K., Greenfield, P., Lyn, H., & Savage-Rumbaugh, S. (2014). Gestural and symbolic development among apes and humans: Support for a multimodal theory of language evolution. Frontiers in Psychology, 5, 1228. https://doi.org/10.3389/fpsyg.2014.01228

Gillespie-Lynch, K., Greenfield, P. M., Feng, Y., Savage-Rumbaugh, S., & Lyn, H. (2013). A cross-species study of gesture and its role in symbolic development: Implications for the gestural theory of language evolution. Frontiers in Psychology, 4, 160. https://doi.org/10.3389/fpsyg.2013.00160

Goodwyn, S. W., Acredolo, L. P., & Brown, C. A. (2000). Impact of symbolic gesturing on early language development. Journal of Nonverbal Behavior, 24(2), 81–103. https://doi.org/10.1023/A:1006653828895

Hardus, M. E., Lameira, A. R., Van Schaik, C. P., & Wich, S. A. (2009). Tool use in wild orangutans modifies sound production: A functionally deceptive innovation? Proceedings of the Royal Society B: Biological Sciences, 276(1673), 3689–3694. http://doi.org/10.1098/rspb.2009.1027

Hewes, G. W. (1973). Primate communication and the gestural origin of language. Current Anthropology, 14(1-2), 5–24. https://doi.org/10.1086/201401

Hopkins, W. D., Taglialatela, J. P., & Leavens, D. A. (2007). Chimpanzees differentially produce novel vocalizations to capture the attention of a human. Animal Behaviour, 73(2), 281–286. https://doi.org/10.1016/i.anbehav.2006.08.004

Iverson, J. M., Capirci, O., & Caselli, M. C. (1994). From communication to language in two modalities. Cognitive development, 9(1), 23–43. https://doi.org/10.1016/0885-2014(94)90018-3

Iverson, J. M., & Goldin-Meadow, S. (2005). Gesture paves the way for language development. Psychological Science, 16(5), 367–371. http://doi.org/10.1111/i.0956-7976.2005.01542.x

Iverson, J. M., & Wozniak, R. H. (2016). Transitions to intentional and symbolic communication in typical development and in autism spectrum disorder. In D. Keen, H. Meadan, N. C. Brady, & J. W. Halle (Eds.), Prelinguistic and minimally verbal communicators on the autism spectrum (p. 51–72). Springer Science + Business Media. https://doi.org/10.1007/978-981-10-0713-24

Iyer, S. N., & Oller, D. K. (2008). Prelinguistic vocal development in infants with typical hearing and infants with severe-to-profound hearing loss. The Volta Review, 108(2), 115.

Iyer, S. N., Denson, H., Lazar, N., & Oller, D. K. (2016). Volubility of the human infant: Effects of parental interaction (or lack of it). Clinical Linguistics& Phonetics, 30(6), 470–488. http://doi.org/10.3109/02699206.2016.1147082

Jhang, Y., & Oller, D. K. (2017). Emergence of functional flexibility in infant vocalizations of the first 3 months. Frontiers in Psychology, 8, 300. https://doi.org/10.3389/fpsyg.2017.00300

Jhang, Y., Franklin, B., Ramsdell-Hudock, H. L., Oller, D. K. (2017). Differing roles of the face and voice in early human communication: Roots of language in multimodal expression. Frontiers in Communication, 2(10), 1–12. https://doi.org/10.3389/fcomm.2017.00010

Kendon, A. (2017). Reflections on the “gesture-first” hypothesis of language origins. Psychonomic Bulletin& Review, 24(1), 163–170. https://doi.org/10.3758/s13423-016-1117-3

Koopsman-van Beinum, F. J., & Van der Stelt, J. M. (1986). Early stages in the development of Speech movements. In B. Lindblom, & R. Zetterstrom (Eds.), Precursors of early speech (p. 37–50). Wenner-Gren Center International Symposium Series.

Lameira, A. R., Hardus, M. E., Mielke, A., Wich, S. A., & Shumaker, R. W. (2016). Vocal fold control beyond the species-specific repertoire in an orangutan. Scientific Reports, 6(1), 1–10.

Lameira, A. R. (2017). Bidding evidence for primate vocal learning and the cultural substrates for speech evolution. Neuroscience andBiobehavioralReviews, 83, 429–439. https://doi.org/10.1016/j.neubiorev.2017.09.021

Lynch, M. P., Oller, D. K., Steffens, M. L., Levine, S. L., Basinger, D. L., Umbel, V. (1995). Onset of speech-like vocalizations in infants with down syndrome. American Journal on Mental Retardation, 100(1), 68–86.

Liszkowski, U., Brown, P., Callaghan, T., Takada, A., & De Vos, C. (2012). A prelinguistic gestural universal of human communication. Cognitive Science, 36(4), 698–713. http://doi.org/10.1111/j.1551-6709.2011.01228.x.

Locke, J. L. (1993). The child’s path to spoken language. Harvard University Press.

Long, H. L., Oller, D. K., & Bowman, D. A. (2019). Reliability of listener judgements on infant vocal imitation. Frontiers in Psychology, 10, 1340. http://doi.org/10.3389/fpsyg.2019.01340

Long, H. L., Bowman, D. D., Yoo, H., Burkhardt-Reed, M. M., Bene, E. R., & Oller, D. K. (2020). Social and endogenous infant vocalizations. PloS One, 15(8), e0224956. http://doi.org/10.1371/journal.pone.0224956

Milenkovic P. (2010). TF32 [Computer Program]. Department of Electrical and Computer Engineering, University of Wisconsin, Madison.

Moulin-Frier, C., Nguyen, S. M., & Oudeyer, P. Y. (2014). Self-organization of early vocal development in infants and machines: The role of intrinsic motivation. Frontiers in Psychology, 4, 1006. https://doi.org/10.3389/fpsyg.2013.01006

Müller, G. B., & Newman, S. A. (2003). Origination of organismal form: Beyond the gene in developmental and evolutionary biology. MIT Press.

Nathani, S., Ertmer, D., & Stark, R. (2006). Assessing vocal development in infants and toddlers. Clinical Linguistics and Phonetics, 20(5), 351–369. http://doi.org/10.1080/02699200500211451

Newman, S. A. (2016). Origination, variation, and conservation of animal body plan development. Cell Biology and Molecular Medicine Reviews, 2, 130–162. https://doi.org/10.1002/3527600906.mcb.200400164.pub2

Oller, D. K. (2000). The emergence of the capacity for speech. Lawrence Erlbaum Associates.

Oller, D. K., Buder, E.H., Ramsdell, H.L., Warlaumont, A.S., Chorna, L.B., & Bakeman, R. (2013). Functional flexibility of infant vocalization and the emergence of language. Proceedings of the National Academy of Sciences, 110(16), 6318–6323. http://doi.org/10.1073/pnas.1300337110

Oller, D. K., Griebel, U., Iyer, S. N., Jhang, Y., Warlaumont, A. S., Dale, R., & Call, J. (2019). Language origins viewed in spontaneous and interactive vocal rates of human and bonobo infants. Frontiers in Psychology, 10, 729. http://doi.org/10.3389/fpsyg.2019.00729

Oller, D. K., Caskey, M., Yoo, H., Bene, E. R., Jhang, Y., Lee, C. C., Bowman, D. D., Long, H. L., Buder, E. H., & Vohr, B. (2019). Preterm and full term infant vocalization and the origin of language. Scientific Reports, 9(1), 1–10. http://doi.org/10.1038/s41598-019-51352-0

Orr, E. (2018). Beyond the pre-communicative medium: A cross-behavioral prospective study on the role of gesture in language and play development. Infant Behavior and Development, 52, 66–75. http://doi.org/10.1016/i.infbeh.2018.05.007

Oudeyer, P. Y. (2005). The self-organization of speech sounds. Journal of Theoretical Biology, 233(3), 435–449. http://doi.org/10.1016/i.itbi.2004.10.025

Oudeyer, P. Y., & Kaplan, F. (2006). Discovering communication. Connection Science, 18(2), 189–206. https://doi.org/10.1080/09540090600768567

Patterson, F. G., & Cohn, R. H. (1990). Language acquisition by a lowland gorilla: Koko’s first ten years of vocabulary development. Word, 41(2), 97–143. https://doi.org/10.1080/00437956.1990.11435816

Pika, S., Liebal, K., Call, J., & Tomasello, M. (2005). Gestural communication of apes. Gesture, 5(1-2), 41–56. https://doi.org/10.1075/gest.5.1.05pik

Pollick, A. S., & De Waal, F. B. (2007). Ape gestures and language evolution. Proceedings of the National Academy of Sciences, 104(19), 8184–8189. http://doi.org/10.1073/pnas.0702624104

Riede, T., Bronson, E., Hatzikirou, H., Zuberbühler, K. (2005) Vocal production mechanisms in a non-human primate: morphological data and a model. Journal of Human Evolution, 48, 85–96. http://doi.org/10.1016/i.ihevol.2004.10.002

Rivas, E. (2005). Recent use of signs by chimpanzees (pan troglodytes) in interactions with humans. Journal of Comparative Psychology, 119(4), 404. https://doi.org/10.1037/0735-7036.119.4.404

Silva Lima, E. D., & Cruz-Santos, A. (2012). Acquisition of gestures in prelinguistic communication: A theoretical approach. Revista da Sociedade Brasileira de Fonoaudiologia, 17(4). https://doi.org/10.1590/S1516-80342012000400022

Sterelny, K. (2012). Language, gesture, skill: the co-evolutionary foundations of language. Philosophical Transactions of the Royal Society B: Biological Sciences, 367(1599), 2141–2151. https://doi.org/10.1098/rstb.2012.0116

Soltis, J. (2004). The signal functions of early infant crying. Behavioral and Brain Sciences, 27(4), 443–458. https://doi.org/10.1017/S0140525X0400010X

Stark, R. E. (1980). Stages of speech development in the first year of life. In G. Yeni-Komshian, J. Kavanaugh, & C. Ferguson (Eds.), Child Phonology (p. 73–90). Academic Press.

Tomasello, M., & Zuberbühler, K. (2002). Primate vocal and gestural communication. In M. Bekoff, C. Allen, & G. M. Burghardt (Eds.), The cognitive animal: Empirical and theoretical perspectives on animal cognition (p. 293–299). MIT Press.

Tomasello, M., Carpenter, M., & Liszkowski, U. (2007). A new look at infant pointing. Child Development, 78(3), 705–722. https://doi.org/10.1111/j.1467-8624.2007.01025.x

Tomasello, M. (2010). Origins of human communication. MIT press.

Volterra, V., Caselli, M. C., Capirci, O., & Pizzuto, E. (2005). Gesture and the emergence and development of language. In M. Tomasello & D. I. Slobin (Eds.), Beyond nature-nurture: Essays in honor of Elizabeth Bates (p. 3–40). Lawrence Erlbaum Associates.

Wich, S. A., Swartz, K. B., Hardus, M. E., Lameira, A. R., Stromberg, E., & Shumaker, R. W. (2009). A case of spontaneous acquisition of a human sound by an orangutan. Primates, 50(1), 56–64. https://doi.org/10.1007/s10329-008-0117-y

Wu, Z., & Gros-Louis, J. (2015). Caregivers provide more labeling responses to infants’ pointing than to infants’ object-directed vocalizations. Journal of Child Language, 42(3), 538–561. https://doi.org/10.1017/S030500091400022

